# Long Range Order and Short Range Disorder in *Saccharomyces cerevisiae* Biofilm

**DOI:** 10.1101/451062

**Authors:** Vincent Piras, Adam Chiow, Kumar Selvarajoo

## Abstract

Biofilm, a colony forming cooperative response of microorganisms under environmental stress, is a major concern for food safety, water safety and drug resistance. Most current works focus on controlling biofilm growth by targeting single genes. Here, we investigated transcriptome-wide expressions of the biofilm yeast *Saccharomyces cerevisiae* in wildtype, and 6 previously identified biofilm regulating overexpression strains (*DIG1, SAN1, TOS8, ROF1, SFL1, HEK2*). When tested across various statistical distributions, all transcriptome-wide data fitted well with lognormal distribution above TPM value of 5. Using this threshold as a low expression filter, Pearson auto-and cross-correlation reveal a strong transcriptome-wide invariance among all genotypes, which is also reflected by the random selection of 50 gene expressions. Focusing on the 50 highly expressed genes, however, they differ significantly between the genotypes. Principal components analysis (PCA) shows global similarity between *DIG1, SAN1, ROF1, SFL1* and *HEK2.* Thus, although single overexpression strains may show significant favourable local and acute expression changes (short range disorder), the almost unperturbed global and collective structure between the genotypes indicate gradual adaptive response converging to original stable biofilm states (long range order). Hierarchical clustering and Gene Ontology show 11 groups of local (e.g. mitochondria processes, amine & nucleotide metabolic processes) and 6 groups of global (e.g. transcription, translation & cell cycle) processes for all genotypes. These data indicate that there is a strong global regulatory structure that keeps the overall biofilm stable in all investigated strains.

## 1. Introduction

Biofilm is a major area of research in food, drug and dental industries, due to their persistence, emergent and adaptive responses [1]. Although biofilm can have positive attributes, most situations focus on the negative consequences. Previous studies have mainly focused on the extracellular polymeric substance formation or quorum sensing mechanisms on low throughput platforms. Despite the progress made in understanding biofilm components and constituents, till-today, there is no key solution in controlling its progression. Thus, there is a general need to venture into high-throughput methodologies to investigate further on the complex collective responses of biofilm.

Transcriptome-wide expression studies using RNA-Seq techniques has gained tremendous momentum in the last decade. The successes have been mainly due to deeper sequencing and wider dynamic ranges compared to its predecessors such as the microarray [2]. An overwhelming majority of works, however, is investigating a limited subset of genes with higher expression levels or fold changes to infer differential regulation, functional pathways or regulatory networks using common statistical approaches. This is often with the belief that lower expression transcripts are marred with technical/operator induced noise or insignificant fold changing genes are not important. Although this is a necessary concern when we analyse and compare individual gene expressions, on a global transcriptome-wide scale, any arbitrary threshold cut-off may have negative outcomes. For example, the popular power-law distribution is only observed when we have a large number of entities within the datasets [3]. Taking a subset of a dataset can lead to the disappearance of the statistical structure [4]. Furthermore, Information theory proposes that the expression level of an entity, such as a gene, carry no real meaning without context, or its probability [5–6]. Thus, analysing high throughput experimental data should not overly focus on the expression values alone (i.e. analysing only highly expressed or significant fold changing data), but look out for the principles used in information theory [7–9].

Living cells are complex, dynamical and dissipative systems considered to be in a state that is far from equilibrium [10, 11]. In other words, living cells are dynamically exchanging matter for their survival, and are able to evolve spontaneously (say, under any external perturbation) towards a critical point, without fine-tuning system parameters, for a phase transition [12, 13]. The formation of biofilm by certain microorganisms, such as the bacteria *E. coli* and yeast, is a classic example [14, 15]. Such phase transformation is known to break the symmetry of the system leaving it invariant, or in a collective mode [16].

Previously, we have used information probability concepts and have shown for diverse mammalian cells, that pair-wise transcriptome-wide expressions, with several orders of magnitude difference between their expression levels, are very high (>0.98) [17–22]. The strong invariance, across the large expression variability between genes, is a signature of order parameter organising the entire gene regulatory network imposed by the presence of a common attractor correspondent to the cell type [23–24].

In this paper, to investigate the yeast *Saccharomyces cerevisiae* biofilm phase transition, we adopted information theory related statistical approaches to analyse the transcriptome-wide RNA-seq expressions in wildtype and 6 overexpressed gene mutants (*DIG1, SAN1, TOS8, ROF1, SFL1, HEK2*) [25, 26]. Our analyses include scatter plots, distribution fitting, Pearson correlation, principal components analysis, hierarchical clustering and gene ontology (GO) enrichment.

## 2. Results

We analysed the whole transcriptome of wildtype and 6 overexpressed strains (*DIG1, SAN1, TOS8, ROF1, SFL1, HEK2*), with 4 replicates for each condition, of biofilm modulators in the yeast *Saccharomyces cerevisiae* strain F45, adapted from Cromie *et al.* [25, 26]. The selected genes are known to regulate the fluffy colony morphology of the yeast strain studied. There was a total of 11,236 unique read counts (genes) with non-zero expression in the datasets across all genotypes (Materials & Methods). We next performed normalisation using TPM method [27].

### 2.1. Statistical frequency distributions show lognormal as most probable

It has been shown on several occasions by others, and also by us, that gene expression distributions follow power-law or lognormal distributions due to their scale-free network organisation or complex transcriptional regulatory mechanisms [2, 29–31]. Hence, we tested the frequency distribution of gene expressions in all genotypes (See Fig. 1 and Fig. S1) using 5 statistical distribution fits: i) log-normal, ii) log-logistic, iii) Pareto (power law), iv) Burr, and v) double Pareto lognormal (DPLN) (Materials & Methods). We observed that, in all datasets, the expression distributions fitted all 5 tested distributions above TPM > 1. The log-normal has a slight advantage for the higher expressed genes (TPM > 5), while the DPLN fitted better at the lower expressed genes (TPM < 5).

**Fig. 1.**
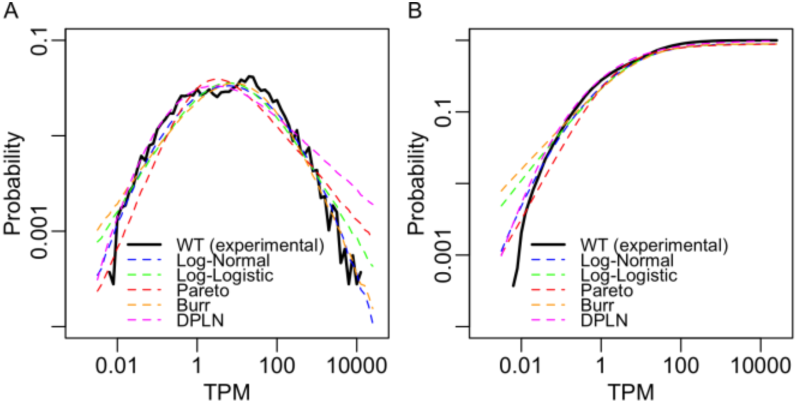
Statistical distribution fitting. Gene expression distributions (in log scale) against **(A)** probability density function (PDF) and **(B)** cumulative density function (CDF). Blue, green, red, orange, and magenta curves indicate log-normal, log-logistic, power-law (Pareto), Burr and double Pareto log-normal fitting (DPLN) respectively. The thick black curve corresponds to the experimental distribution in the wildtype (WT) condition (average of 4 replicates). Parameters of the fitted distributions are summarised in Table S1.

This result suggests that the two tails of the gene expression distributions might be due to two different regulations [32]. The upper tail corresponds to higher gene expressions, from transcripts produced through more constitutive gene expression mechanisms, less prone to biological variations or to technical noise. The lower tail corresponds to lower gene expressions, arising from rare transcripts generated from highly stochastic (noisy) transcriptional events, and the measurement of these low expressions is more prone to technical errors [31,33]. It will, therefore, be important to carefully determine a threshold to discriminate between genes that carry important information on the biological response from noisy data.

From the statistical structure, the threshold is at about TPM = 5, and we will scrutinise this threshold on the next section using correlation.

### 2.2. Strong transcriptome-wide invariance between biological replicates and different overexpression conditions

We computed pair-wise Pearson correlation, *r*, for the whole transcriptome (*N* = 11,236) between the replicates of each genotype (See Fig. S2). The result shows that the whole transcriptome correlation between each replicate is very close to 1 (min = 0.979, mean > 0.987, See Fig. 2A, left panels) for all genotypes. Such strong transcriptome-wide correlation invariance has also been observed for other cell types [17–22]. Next, comparing cross-correlations, *r*_*c*_, (correlations between genotypes), although the values are lower, they also remain near unity albeit slightly lower (mean between 0.945 and 0.972, See Fig. 2A, right panels). The data suggest that the global collective regulation is keeping the transcriptome-wide correlation high, despite the acute expression changes of a limited gene subset between the genotypes.

**Fig. 2.**
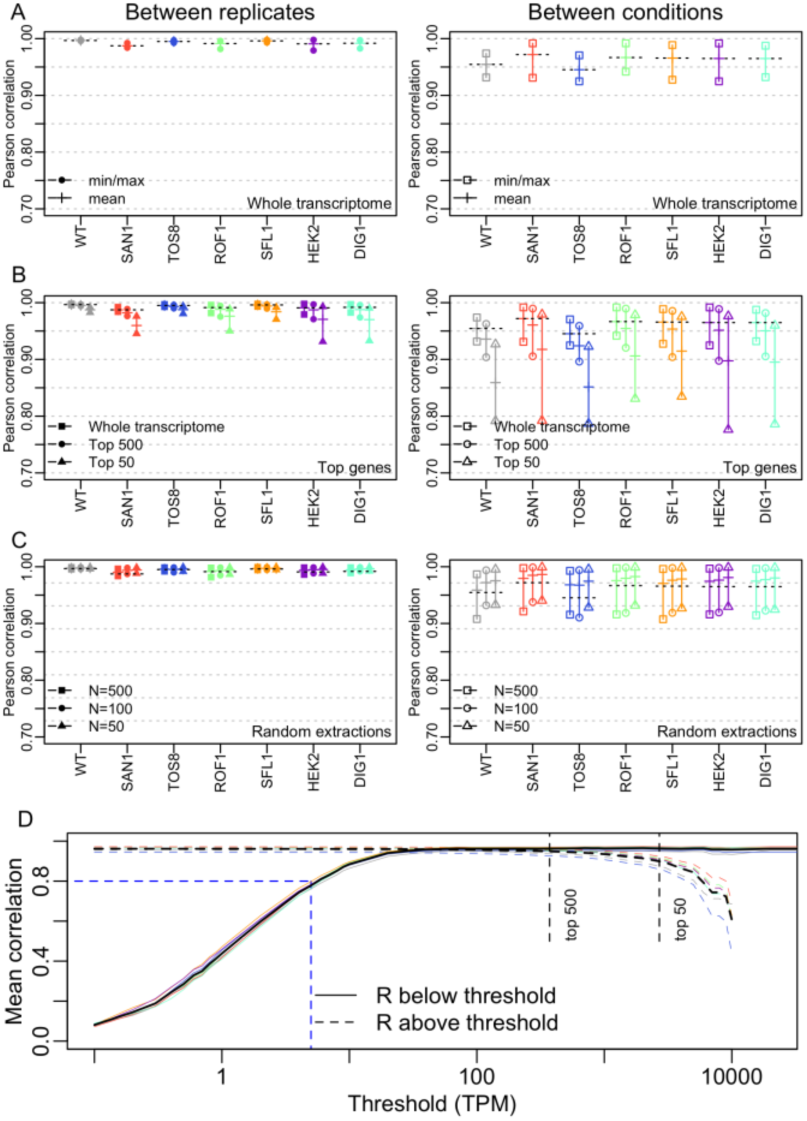
Correlation Plots. Pearson (auto-and cross-) correlations for **(A)** whole transcriptome (*N* = 11,236), **(B)** top 500 and top 50 most expressed genes in each condition and **(C)** random gene extractions with sample sizes *n* = 500, 100 and 50. Filled shapes (on left panels): range (min, max) of values taken by Pearson correlations between replicates (*r*). Empty shapes (on right panels): range of Pearson correlations against other conditions (*r*_*c*_), for each condition. Dashes: mean correlation values in the range. Dotted lines (repeated in every plot): whole transcriptome average for each condition. Note: cross-correlation values (min, max and mean) were obtained from all between-sample combinations (i.e., each condition’s 4 replicates against the 4 replicates of each of the 6 other conditions). **(D)** Mean *r*_*c*_ when selecting transcriptome size above (dashed) and below (solid line) an expression threshold (TPM, x-axis). For TPM < 5 (dashed blue line), mean *r*_*c*_ < 0.8. Individual replicates are indicated by thin coloured dashed lines.

To observe the effect of highly expressed transcripts on the correlation values, *r* and *r*_*c*_ values were evaluated for the top ranked 50 and 500 highest expressed genes (See Fig. 2B). Notably, the *r-*values remained very high (mean between 0.960-0.988). However, the *r*_*c*_ values were noticeably lowered. For example, the mean *r*_*c*_ between WT and other conditions for the 50 highest genes expressed is 0.859, while it is 0.936 for 500 highest expressed genes and 0.955 transcriptome-wide. Note that removing either 500 highest or lowest expressed genes from the transcriptome had little effect on *r* and *r*_*c*_ values compared with whole transcriptome (See Fig. S3A and B).

Next, we evaluated the correlations for selections of 100 expressed genes with mean expression values around 100 TPM (highly expressed, see expression distributions, See Fig. 1), 5 TPM (in the middle of the distribution), and 1 TPM (lowly expressed). For genes with TPM∼100, *r* and *r*_*c*_ values remained very high, close to correlations obtained for whole transcriptome, however, both *r* and *r*_*c*_ values were significantly decreased for lower gene expressions: for genes with TPM ∼ 5, *r* and *r*_*c*_ values still range at positive values around 0.5, while for TPM∼1, the correlations were close to 0. (See Fig. S3C and D). This result confirms that genes with TPM < 5 carry high noise that destroys the correlation structure.

Finally, to test whether sample size impacts correlation values, we tested various sizes for random genes selections (Materials & Methods) for each genotype 100 times and computed their *r* values (See Fig. 2C). Notably, both *r* and *r*_*c*_ values are significantly higher than the highest expressed genes, and similar to those of whole transcriptome values for all genotypes. Thus, the global or transcriptome-wide statistical structure is also revealed by the random selection of genes rather than choosing the highest expressed ones. In other words, random selection of genes shows the fractal nature of transcriptome-wide gene expressions.

Overall, these data show that the highest or lowest expressed genes between different genotypes are differentially regulated causing lowering of their correlation values (See Fig. 2D): highly expressed genes show lower correlation because of their large variance in expression values (larger range of activation). In contrast, the decrease in correlations for lower expressed genes is due to non-informative noise. These data suggest that on a global scale, the technical and biological noise for the gene expressions is not sufficient enough to destroy the statistical structure of all genotypes and replicates investigated for most of the transcriptome (TPM > 5, *N* = 6,328). This result agrees with our previous work on neutrophil gene expression study in that the lowly expressed transcripts may not be wasteful but could play important roles in cell function [24]. It is, therefore, important to check the statistical structure before arbitrarily discarding a large portion of lowly expressed genes assuming that they carry no information or simply noisy. Hence, based on both statistical structure (distributions) and correlation, we chose to retain 6,328 genes with TPM > 5 for further global analysis.

### 2.3. Principal Component Analysis and statistical clustering reveal similarity/diversity between overexpressed conditions

We evaluated the principal components (PC) for different sample sizes of sorted genes from the highest to lowest expression values (*N* = 10, 30, 50, 100, 150, 200, 300, 500 and 6,328, i.e. TPM > 5, See Fig. 3A-C). PCs 1 to 3 constituted more than 76% variance (in whole informative transcriptome, *N* = 6,328, See Fig. S4A), and we plotted them for each genotype for the different *N* sizes (See Fig. S4B). Notably, the PC loading scores (position) of each genotype became fixed when *N* ≥ 100 (See Fig. S4C). That is, a minimum sample size of 100 highest expressed genes is sufficient for the convergence of PCs with the whole transcriptome data. If *N* < 100, the PC values became more variable and overlapped with other genotypes (See Fig. S4B-C)

**Fig. 3.**
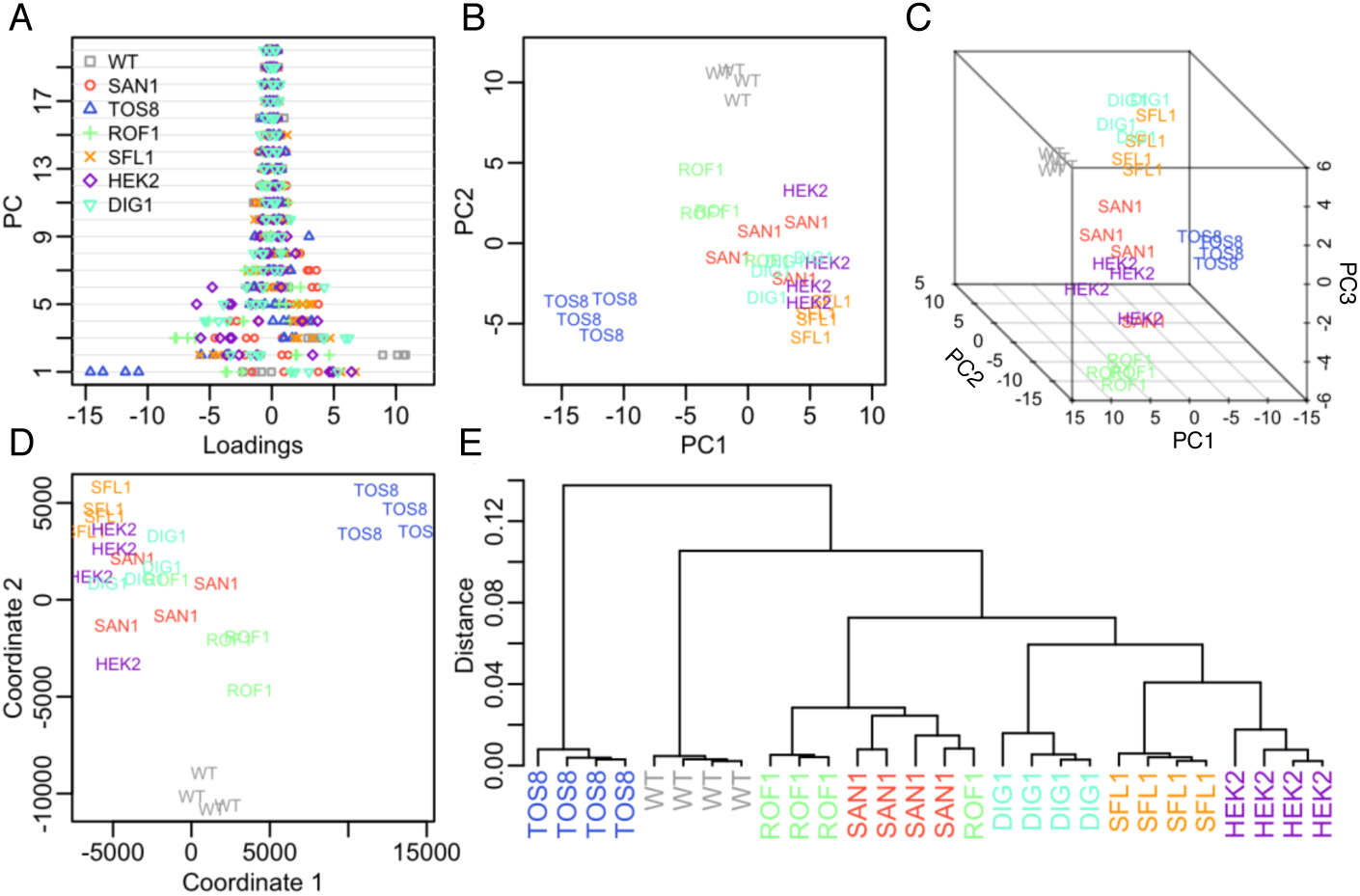
Principal component and clustering. **(A)** PCA loadings for the first 20 components. Note for PCA1 the clustering of the 4 TOS8 condition replicates (blue triangles) at negative values, for PCA2 the clustering of the 4 WT replicates at positive values (grey squares) and for PCA3 the clustering of the 4 ROF1 replicates at negative values (green crosses). **(B)** 2D projection of the 2 first PCs. **(C)** 3D representation of the 3 first PCs. **(D)** Multidimensional scaling representation of the 28 samples using Euclidean distance. **(E)** Hierarchical clustering using Pearson correlation as distance metric and Ward method for tree construction.

Looking closer at PCA loadings (See Fig. 3A, left), we observe PC1 is able to discriminate replicates in the TOS8 overexpression condition with no overlap with other conditions, while extreme loading scores for PC2 are obtained for the replicates of the wildtype condition, and PC3 extreme scores are for the ROF1 condition. Other PC vectors, in contrast, show low specificity with high overlap between replicates of different conditions. When plotting PC1 vs. PC2 and PC3, we also observe clustering and seclusion of the wildtype replicates, as well for TOS8 replicates (See Fig. 3A, middle and right). Replicates of other conditions also tend to cluster together, however, with more overlap with other conditions. This is also confirmed by multidimensional scaling (MDS) over Euclidean distances between replicates and conditions (See Fig. 3B), which shows that the largest distances between conditions are between wildtype, TOS8 and SFL1. Together, PCA and MDS indicate that wildtype and TOS8 conditions have the most distinct variance (response) while other conditions are closer in terms of transcriptome-wide response. Hierarchical clustering of the expression matrix with (TPM > 5, See Fig. 3C), based on Pearson correlation as distance metric, corroborates these results, and indicated 4 major clusters: i) all 4 wildtype replicates, ii) TOS8, iii) SFL1/DIG1 and iv) ROF1/HEK2/SAN1. This result is also reflected from the *r*_*c*_ values (see See Fig. 2A, right panel, mean values of *r*_*c*_ for wildtype and TOS8 are the lowest).

### 2.4. Characterisation of differentially and commonly expressed genes between overexpressed conditions

It is noteworthy to identify differentially expressed genes between wildtype and the overexpressed conditions. One way to identify differentially expressed genes is through standardisation of gene expression values (Z-scores) [21]. In brief, the expression value (*x*_*ij*_) of each gene (*i*) in each experiment (replicate or condition, *j*) is normalised to the mean expression values across all experiments (*µ*_*i*_) and scaled by the standard deviation of expression values across all experiments (*σi*), that is: *Z*_*ij*_ *=* (*x*_*ij*_ *– µ*_*i*_)/ *σi*. We performed a log_10_ transformation of the data prior to normalisation to minimise outlier effects, and averaged the Z-scores of all replicates (28 samples x 6,328 genes) for each condition, to obtain an averaged normalised matrix (7 conditions x 6,328 genes).

In order to interpret Z-values, it is important to verify the normality of their distribution (See Fig. S5A): performing a QQ-plot showed most of the distribution approximates the standard normal distribution (mean = 0, s.d. = 1), except for |*Z*| > 2, indicating significant outliers (corresponding *p* < 0.05 assuming normal distribution, See Fig. S5B). As verification, we checked the expression values and Z-scores of the different overexpressed genes in all conditions and confirmed: i) all 6 overexpressed genes show TPM > 5 in all conditions and replicates (See Fig. S5C), ii) all 6 overexpressed genes are significantly overexpressed in their respective overexpression condition, with Z-scores > 2 (See Fig. S5D).

Hence, we identified differentially expressed genes with TPM > 5 and |*Z*| > 2 in at least one experiment, and found, out of the 6,328 genes, only 296 genes, corresponded to the acute transcriptome response to perturbation, that is, genes whose expression change is of similar order to the perturbed (overexpressed) gene (SAN1, TOS8, etc.). This data, therefore, suggest that only a small proportion of the transcriptome is directly regulated by the overexpressed genes, and a large majority are weakly or marginally affected.

### 2.5. Characterisation of the local acute and global collective transcriptome responses in the different overexpressed conditions

To characterise the genes that are dissimilar between wildtype and other conditions and their functions, we first performed hierarchical clustering on the Z-score matrix of the 296 differentially expressed genes that form the acute response to perturbation (TPM > 5 and |*Z*| > 2, see Fig. S7). As proof of principle, we confirmed the 7 overexpressed genes (SAN1, TOS8, HEK2, SFL1, DIG1 and ROF1) were all present in the list. We obtained 32 super-clusters that were then iteratively merged according to their similarity by minimising the correlation between clusters and maximising correlation within clusters (See Fig. S6A). After the procedure, we obtained 11 distinct clusters (See Fig. S6B and C), of which, 7 consist of upregulated genes in each condition, and 4 consist of down-regulated genes in wildtype, DIG1, SFL1 and TOS8 conditions. Note that down-regulated genes in wildtype indicate up-regulated genes in all overexpression experiments (and vice-versa).

The two largest clusters correspond to up-and down-regulated genes in wildtype compared to all other conditions (up-regulated *n* = 69 genes, cluster Ac/C2 – <u>Ac</u>ute <u>C</u>ommon <u>2</u>and C; down-regulated *n* = 128, cluster Ac/C1, See Fig. 4A). In other words, from the 296 differentially expressed genes, 197 were common between all overexpression conditions (67%), indicating a core set of genes that defines a non-specific response. Performing gene ontology (GO) using AMIGO2 [34] showed the up-regulated genes of the non-specific acute response are involved in various mitochondrial processes, in particular translation, while the down-regulated genes play a role in the nucleotide and amine metabolic processes (Table S2). On top of this, TOS8 specifically regulated 80 genes (21 up-and 59 down-regulated, clusters Ac/S3 and Ac/S8 – <u>Ac</u>ute <u>S</u>pecific <u>3</u> and <u>8</u>, 27%). Notably, genes involved in the fungal-type vacuole membrane were down-regulated by TOS8. In other conditions, very few other genes were specifically differentially expressed.

**Fig. 4.**
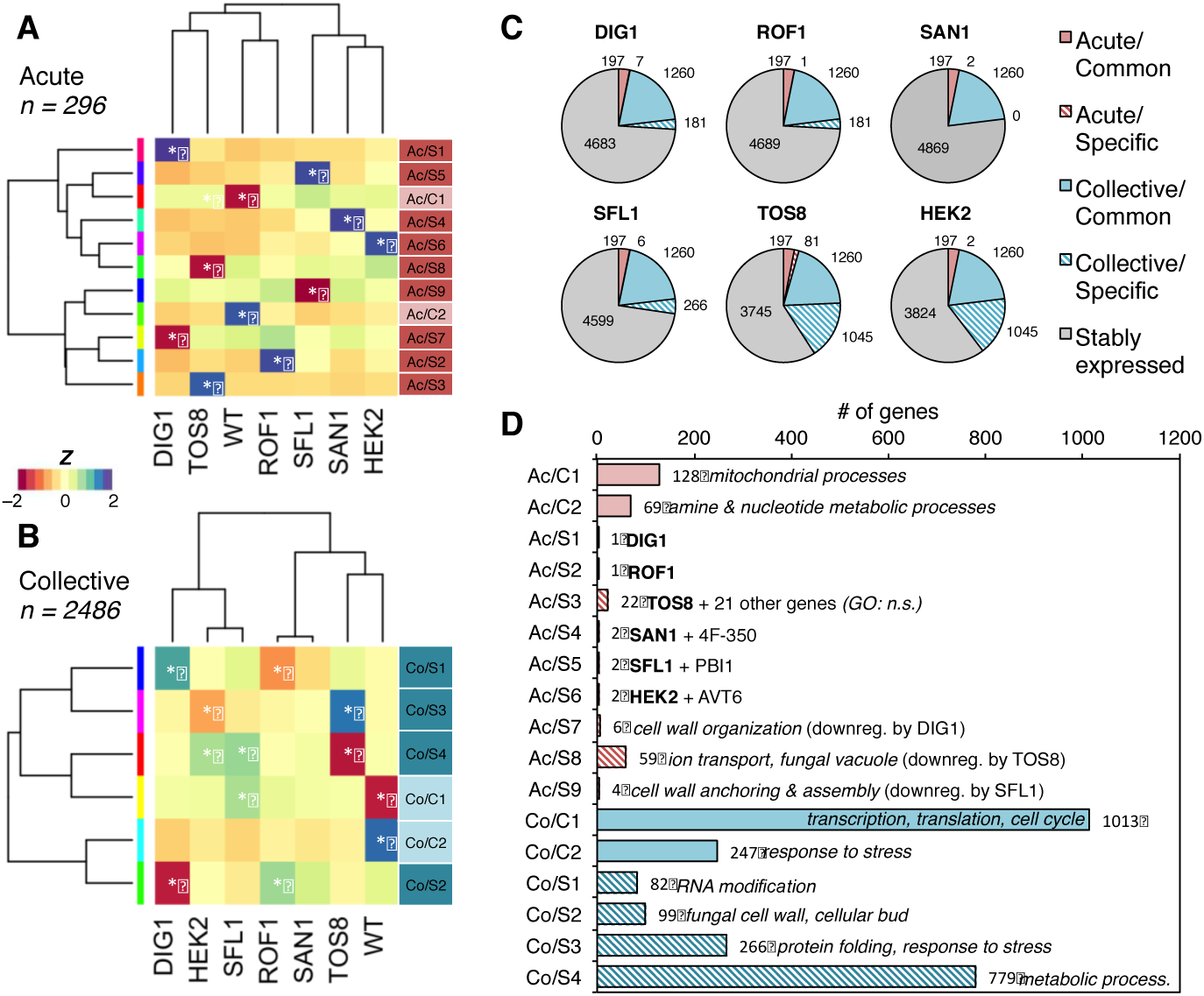
Gene expression clustering. Heatmaps of average Z-values of the 11 clusters (9 condition-specific: Ac/S*x*, and 2 common to all conditions: Ac/C*x*) obtained for **(A)** the acute response (296 genes with TPM > 5 and |*Z*| ≥ 2), and **(B)** the 6 clusters (4 condition-specific: Co/S*x* and 2 common: Co/C*x*) of the collective response (2,486 genes with TPM > 5 and 1.5 ≤ |*Z*| < 2). Positive (blue/green) and negative (red/orange) values correspond to up-and down-regulated clusters respectively. Note up-regulation (blue) in the wildtype (WT) condition indicates down-regulation in all other single gene overexpression condition (and vice-versa). Stars indicate selected clusters with most important gene expression changes for each condition. **(C)** Distribution of the 6,328 expressed genes (TPM > 5 in at least one condition, ∼52% of the detected transcriptome) for each overexpression condition: common categories (in plain colors) contain genes that have significant expression changes in all single-gene overexpression conditions. Specific categories (in striped patterns) are genes with significant expression changes in some conditions. Stably expressed genes (in gray) show no significant expression changes (|*Z*| < 1). Number of genes for each category is indicated. **(D)** Number of genes for each cluster from the acute (Ac, in pink) and collective responses (Co, in light blue). Enriched GO terms are indicated in italic for each cluster, when present (see detailed list in Table S2 and S3).

Other overexpressed conditions also regulated a small number of genes: DIG1 (alone, cluster Ac/S1) down-regulated 6 genes (cluster Ac/S7) and SFL1 regulated 5 other genes (1 up, 4 down, clusters Ac/S5 and Ac/S9). HEK2 upregulated another gene (cluster Ac/S6), and so did SAN1 (cluster Ac/S4). ROF1 did not show any specific acute response (cluster Ac/S2 only contained ROF1 gene). For these clusters, because of the low number of genes, GO could not be performed.

These results indicate that the acute response is very limited to only a small number of genes, and for a large majority, belong to very generic and non-specific biological processes, e.g. to support the transcriptional and translational response to an external or internal stress. Notably, we have previously shown that genes with lower expression changes can also play an important role in shaping collective and coordinated transcriptional responses [18,24]. It is, therefore, interesting to investigate the interplay between the acute and collective responses in the different overexpression conditions.

We, thus, expanded the pool of differentially expressed genes by lowering the Z-values threshold, that is, to include genes with much weaker (yet significant) gene expression changes compared to the acute response, but that collectively respond to a perturbation [18,24]. Based on the determined threshold (see Fig. S8 and Materials and Methods, *Determination of the collective response genes*), we could include 2,486 additional genes with 1.5 < |*Z*| ≤ 2 that form the collective global response (see Fig. S7). We again performed hierarchical clustering of the Z-values resulting in 48-super clusters (See Fig. S6D), and finally obtained 6 distinct expression profiles (See Fig. 4B and Fig. S5E and F). Similarly, to the acute response, we also observe a large group of genes that are commonly up-or down regulated in all conditions (1260 genes, clusters Co/C2 and Co/C1 – <u>Co</u>llective <u>C</u>ommon <u>1</u> and <u>2</u>, See Fig. 4B and C). These genes belong to diverse biological processes involved in the collective transcriptome aspecific response (transcription, translation, cell cycle, response to stress, etc., Table S3).

We also observe a number of clusters that are specific to certain overexpressed conditions. However, in contrast to the acute response, where we found only one-condition-specific clusters, we see a more intertwined regulation of the clusters between the different conditions (See Fig. 4C). For example, 779 genes that are up-regulated in TOS8 condition (cluster Co/S4 – <u>Co</u>llective <u>S</u>pecific <u>4</u>) and involved in many metabolic processes, are also weakly down-regulated by HEK2 (See Fig. 4B and 4C). Conversely, 266 genes (cluster Co/S3) enriched in processes such as protein folding are down-regulated by TOS8 but upregulated by HEK2 (see Table S3). A similar pattern occurs between DIG1 and ROF1 conditions for 2 smaller clusters (99 genes enriched in fungal cell wall, and 82 genes in RNA modifications, clusters Co/S2 and Co/S1). We also observe that SFL1 does not only down-regulate genes in the collective response but also weakly up-regulate genes that are up-regulated by TOS8. (cluster Co/S3). Finally, the specific collective response seems absent when overexpressing ROF1.

Overall, the analysis indicates that, on top of a very limited acute response (296 genes, of which 197 are non-specific to any overexpression condition), there exist a weak but co-regulated global collective transcriptome response that encompasses a large portion of the transcriptome (2,486 genes, 1260 non-specific) (See Fig. 4C). Most of the differentially expressed genes, either from the acute or collective response, belong to generic biological processes, such as transcriptional, translational, or metabolic processes, that are collectively activated or repressed upon external stress to adapt to diverse environmental changes (Table S2 and S3). Thus, despite each overexpressed condition showing some specific and strong expression changes of a limited genes subset, the global transcriptome structure remains almost unaffected as a vast majority (59%∼76% depending on the condition, general biological processes, see GO terms in Table S4) of the expressed transcriptome is stable (Fig. 4C), and gradually adapts in order to converge to a phenotypic stable state.

## 3. Discussion

Single cell microorganisms are free-living system often yielding uncorrelated (disordered) responses [31–33]. However, when they are exposed to environmental or pathogenic threats, a large number of species are able to aggregate, often within a self-secreted extracellular polymeric substances matrix or through quorum sensing, resulting in a highly ordered structure called biofilm [14].

Self-organized criticality is a general property of dynamical systems, where the overall behaviour shows a critical point leading to a phase transition that will leave the system invariant, or in a collective mode [12]. That is, at the critical point, all differences between individuals will be reduced to follow universal properties leading to long range order. Below the critical point (subcritical), the system will be in disorder [36].

In this paper, we investigated the global properties of the biofilm formed by the *Saccharomyces cerevisiae*. Numerous studies have investigated a number of strains that regulate the growth of the biofilm, however, the conclusions do not point to a clear winner [15, 30]. Here, we adopted a number of statistical metrics to analyse the transcriptome-wide RNA-Seq expressions in wildtype and 6 overexpressed gene conditions (*DIG1, SAN1, TOS8, ROF1, SFL1, HEK2*) of the yeast strain F45.

After accounting for technical noise, we observed log-normal gene distribution (See Fig. 1). Using this statistical structure for expression threshold cut-off, we analysed a total of 6,328 gene transcripts. Pearson correlation reveals strong invariance not only between the same genotype replicates, but also between all genotypes (See Fig. 2). The strong invariance was also recapitulated by random selection of genes. However, when only the highly or lowly expressed genes were compared between genotypes, the correlations were noticeably lower. Thus, we observe two modes of behaviour: a strong global correlation from low to high expression (long ranger order), driven by stably expressed and collectively responding genes and a poorer local correlation at the highest gene expression changes (short range disorder) that carry acute response (see Fig. S7, correlations of acute response genes only vs. transcriptome without acute response genes).

Principal component and clustering (See Fig. 3) showed that TOS8 was the most distinct response from wildtype and any other overexpressed condition. Other conditions were closer to each other in terms of response, while still being quite distinct from wildtype. These simple approaches, thus, reveal the similarity and diversity between the different overexpression and wildtype conditions.

Focusing only on a selected highly expressed genes show short-range disorder. It also revealed that a majority of genes (67% of the acute response, and 51% of the collective response) are commonly regulated between all conditions and GO analysis reveals the distinct functions of the differentially regulated genes (See Fig. 4).

Overall, our analysis provided statistical evidence that biofilms are highly structured transcriptome-wide (not only in their physical appearance), and single mutant/overexpression regulation, such as the ones investigated here, are only able to propagate response locally, that is, to a limited number of genes. Hence, these conditions will likely provide only a transient success in biofilm control and may be subdued in the long run.

These data coincide with the notion of attractor states in biology [24,33,35], where there is a stable equilibrium state towards which a dynamical system tends to converge despite perturbation. Moving out of the attractor state requires the collective action of the entire system parameters, in this case, gene expressions [24,37]. Thus, biofilm control using single gene target may not be a viable option for successful long-term effects. Our data, therefore, ask for a deeper consideration for the understanding of the global statistical structure or organization of dynamic living systems. We, therefore, urge the microbiologist communities to tackle biofilm challenge adopting multidisciplinary approaches that will shed light not only on the acute local effects but also on the collective global regulation, and to identify novel targets for generating long-range disorder.

## 4. Materials and Methods

### 4.1. Datasets

Experimental data was obtained from datasets publicly available on the Gene Expression Omnibus Database (https://www.ncbi.nlm.nih.gov/geo/), under accession numbers GSE98079 and GSE85843 [25,26]. The available FPKM expression data was used and reprocessed into TPM expression units for this study [27], such as the TPM value of the *i*^th^ gene is the ratio between the FPKM value of the *i*^th^ gene and the sum of the FPKM values of all *n* genes, multiplied by a factor for 10^6^:

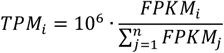

The used data contains 28 samples (4 replicates for 7 conditions). The total transcriptome size is 12,131 genes (distinct SGD IDs), of which 11,236 were detected in at least 1 sample (TPM > 0). 6,328 genes were considered as expressed (TPM ≥ 5) in at least 1 sample, of which 3,546 were considered as non-responsive (|*Z*| < 1), 2,486 from the collective response, and 296 for the acute response.

### 4.2. Statistical Distributions Fitting

Fitting of the distributions of experimental gene expression data was performed using the Maximum-likelihood Fitting method (*fitdistr* of the R MASS package [38] for Pareto, log-normal, log-logistic and Burr distributions. The double Pareto log-normal distribution fitting was performed using a general simulate-annealing approach (GenSA package [39]) to reduce error between experimental PDF curve and the simulated one.

The Pareto (power law) distribution [40] is characterised by 2 parameters, the exponent, *α* that determines the slope of the upper tail of the distribution, and a threshold *x*_*m*_ below which the power law does not apply. The probability density function is defined as, for *x* > *x*_*m*_:

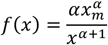

The log-normal distribution is defined for random variable whose logarithm is normally distributed. It is characterised by 2 parameters, *μ* and *σ*, respectively the (log) mean and standard deviation of the underlying normal distribution. Its probability density function (PDF) is defined as:

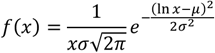

The log-logistic distribution [40] is defined for random variable whose logarithm has a logistic distribution. It is characterised by 2 parameters, *α* and *β*, respectively the scale and shape the underlying logistic distribution. Its PDF is defined as:

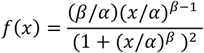

The Burr distribution [40] is one of the generalised log-logistic distributions and is characterised by 3 parameters, *α, β*, and *s*, respectively the 2 shape parameters and the scale parameter. The PDF is defined as:

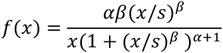

The double Pareto log-normal distribution [41] arises out of a mixture of lognormal distributions shows Pareto power-law behaviour in both tails. It is characterised by 4 parameters, *α, β, τ* and *?*, which stand for the shape parameters (exponents) of the 2 underlying Pareto laws, and the mean and standard deviation of the underlying log-normal distribution. The PDF is defined as:

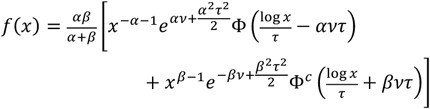

where Φ and Φ^c^ are the cumulative density function (CDF) and complementary CDF of the normal distribution, *N*(0, 1).

### 4.3. Pearson correlation

The Pearson correlation coefficient *r* between two vectors (e.g. transcriptome in two different samples), containing *n* observations (e.g. gene expression values), is obtained by (for large *n*):

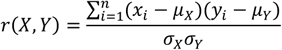

where *x*_*i*_ and *y*_*i*_ are the *i*^*th*^ observation in the vectors ***X*** and ***Y***, respectively, *μ*_*X*_ and *μ* _*Y*_, the average values of each vector, and *σ*_*X*_ and *σ*_*Y*_, the corresponding standard deviations. Pearson correlation measures linear relationship between two vectors, where *r* = 1 if the two vectors are identical, and *r* = 0 if there are no linear relationships between the vectors.

### 4.4. Hierarchical Clustering

Hierarchical clustering builds a hierarchy of clusters using two methods: agglomerative and divisive algorithm. We used the former (Ward’s) where each observation starts in its own cluster, and pairs of clusters are merged moving up the hierarchy. The Ward’s method [42] starts with *n* clusters of size 1 and continues until all the observations are included into one cluster. It begins with the “leaves”, looks for groups of leaves to form “branches” and work its way to reach the “trunk”.

Here, the *n* differentially expressed genes (*n* = 296 for acute response, *n* = 2486 for collective response) were used to form clusters using the normalised gene expression standard scores,

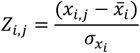

where *x*_*i,t*_ is the expression value (TPM) of the *i*^*th*^ gene at sample *j*, 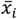 is the mean value in all samples and 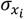 is the standard deviation.

In the first step, *K* clusters are formed (we chose *K* = 32 and *K* = 48 from the initial *n* = 296 and *n* = 2486 genes respectively) by cutting the tree (cutree function in *R*). The next step involves re-clustering the initial *K* clusters into *K*-1 clusters, so the two closest clusters (in terms of distance) are merged. The procedure is repeated until *K* = 1. At each step, we measure the within-cluster correlations (the average correlation among genes that form each cluster), as well as between-cluster correlations (between average profiles of each pair of clusters). We chose the optimal number of clusters, *K*, such that the between-cluster correlations are low, and the within-cluster correlations are high. If *K* is too big, many clusters will have similar profiles between samples (with high between-cluster correlation). Conversely, if *K* is too small, the algorithm will start to merge clusters with heterogeneous gene profiles, and within-clusters correlations will drop.

### 4.5. Multidimensional scaling

Multidimensional scaling (MDS) [43] is a dimensionality reduction approach based on a dissimilarity (distances) matrix between variables (here transcriptomes in different conditions). MDS lays the variables on a 2-Dimensional space, such as the 2D-space Euclidean distances reproduces, with minimum error, the dissimilarity (observed distances) matrix used as input. The metric used for the dissimilarity matrix can vary. Here, we used *n*-D Euclidean distances between *M* transcriptomes as dissimilarity matrix (here, *M* = 28 samples). The distance between two transcriptomes *T*_*j*_ and *T*_*k*_ (e.g. wildtype and SAN1 overexpression conditions) formed by *n* = 6,328 gene expression values (*t*_*j,* 1_,…,*t*_*j*,i_, …, *t*_j,N_) is

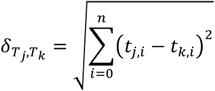

On the 2-D space (with *x-y* coordinates), the reproduced distance between transcriptomes *T*_*1*_ and *T*_*2*_ is defined by

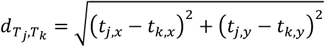

MDS will iteratively rearrange the positions *T*_*j*_ and *T*_*k*_ on the 2D-plane to minimize the following function of a set of *m* transcriptomes

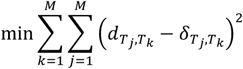

### 4.6. Determination of the collective response genes

To reliably determine from which threshold genes can still be considered as responsive, we first searched for the lowest between-replicate correlation value to determine the maximum correlation shift caused by random variations due to technical noise and biological variability. We found *r* = 0.979 to be the lowest value in our dataset among 42 possible between-replicate correlation values (6 combinations, for 7 conditions), that occurs between 2 different HEK2 replicates. Thus, when comparing two experiments, only a correlation lower than 0.979 would indicate a reliable difference or response between the samples.

We next sorted the genes (with TPM ≥ 5) from high to low absolute *Z*-values. We removed genes one by one, starting from the gene with highest absolute *Z*, and computed the average of cross-correlations (between conditions) for the remaining transcriptome. The average of cross-correlations increases gradually from 0.965 (whole transcriptome that contains all responsive genes), to 0.999 when removing genes with |*Z*| ≥ 1 and above. In particular, removing genes with |*Z*| ≥ 1.5 decreases correlations down to *r* < 0.979. Thus, genes with |*Z*| ≥ 1.5, can reliably be associated with actual expression changes (Fig. S8).

## Supplementary Figures and Tables

**Table S1.** Parameters of the distribution fitting.

|  | Log-normal |  | Log-logistic |  | Pareto |  | Burr |  |  | Double Pareto Log-normal |  |  |  |
| --- | --- | --- | --- | --- | --- | --- | --- | --- | --- | --- | --- | --- | --- |
| | $\mu$ | $\sigma$ | $\beta$ | $\alpha$ | $\alpha$ | $x_m$ | $\alpha$ | $\beta$ | $s$ | $\nu$ | $\tau$ | $\alpha$ | $\beta$ |
| **WT** | 1.887 | 2.459 | 0.697 | 6.808 | 0.495 | 1.71 | 1.629 | 0.607 | 21.261 | 1.394 | -0.226 | 2.599 | 0.254 |
| **SAN1** | 1.913 | 2.455 | 0.695 | 6.979 | 0.49 | 1.709 | 1.673 | 0.603 | 23.365 | 1.394 | 0.062 | 2.157 | 0.132 |
| **TOS8** | 1.785 | 2.55 | 0.67 | 6.217 | 0.458 | 1.275 | 1.855 | 0.568 | 28.266 | 1.538 | 0.176 | 1.742 | 0.023 |
| **ROF1** | 1.885 | 2.493 | 0.684 | 6.87 | 0.476 | 1.55 | 1.89 | 0.579 | 31.653 | 1.433 | 0.001 | 2.095 | 0.073 |
| **SFL1** | 1.906 | 2.502 | 0.683 | 7.007 | 0.475 | 1.579 | 1.873 | 0.578 | 31.764 | 1.437 | -0.107 | 2.301 | 0.06 |
| **HEK2** | 1.838 | 2.521 | 0.68 | 6.562 | 0.475 | 1.486 | 1.833 | 0.578 | 28.297 | 1.502 | 0.088 | 2.036 | 0.09 |
| **DIG1** | 1.906 | 2.476 | 0.687 | 6.874 | 0.474 | 1.543 | 1.649 | 0.597 | 22.613 | 1.49 | 0.075 | 2.269 | 0.157 |

**Table S2.**
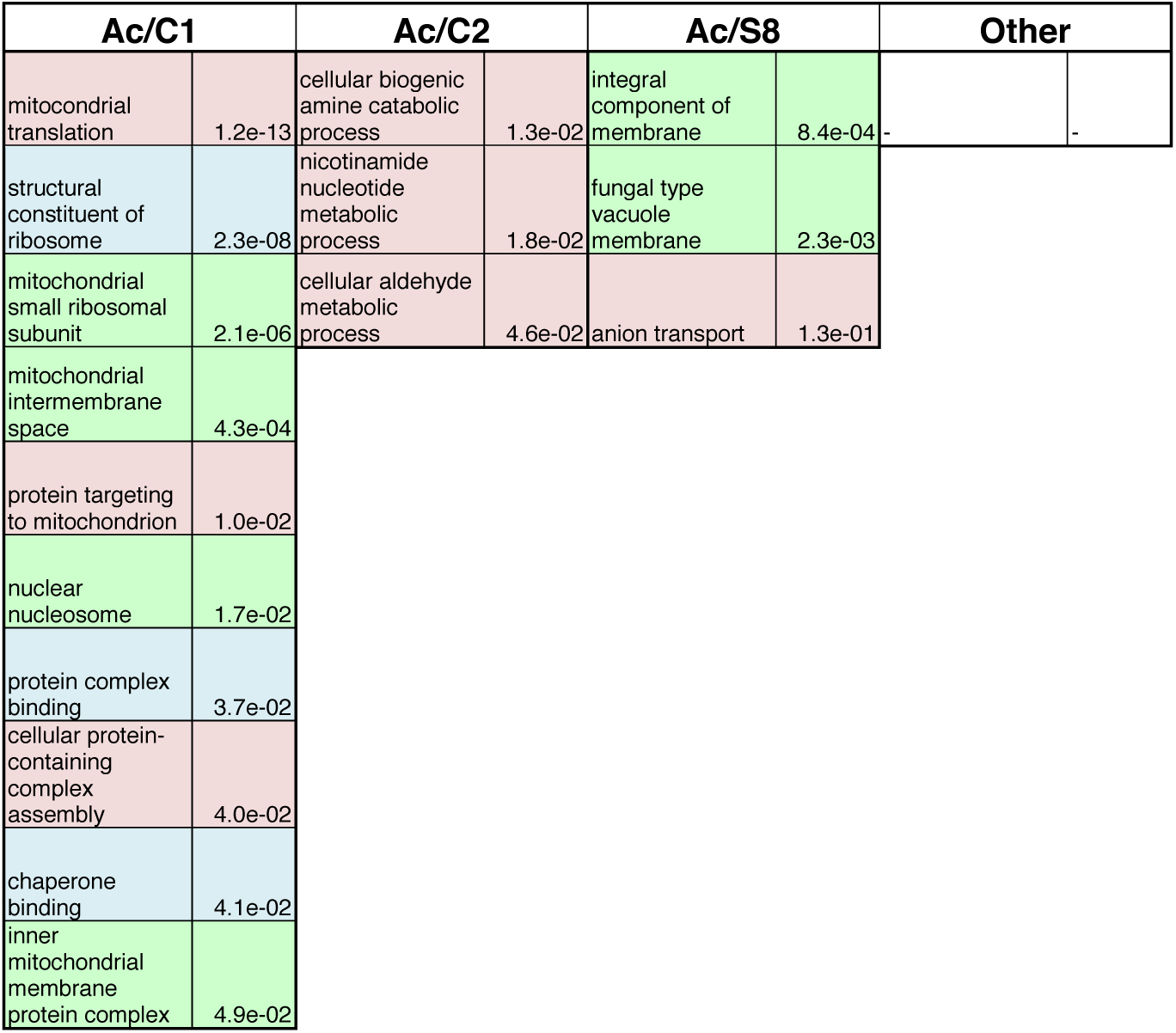
GO processes for acute response clusters. Enriched terms were obtained using AMIGO2. FDR score is reported. Biological process terms are indicated in red background, cellular component terms in green and molecular function terms in blue.

**Table S3.** GO processes for collective response clusters. Enriched terms were obtained using AMIGO2. FDR score is reported. Biological process terms are indicated in pink background, cellular component terms in green and molecular function terms in blue.

| Co/C1 |  |  |  |  |  | Co/C2 |  |
| --- | --- | --- | --- | --- | --- | --- | --- |
| structural constituent of ribosome | 4.07e-15 | microtubule-based process | 8.22e-03 | lagging strand elongation | 2.99e-02 | integral component of membrane | 7.30e-03 |
| cytoplasmic translation | 1.94e-10 | mitochondrial small ribosomal subunit | 9.02e-03 | nucleosome organization | 3.01e-02 | fungal-type vacuole | 1.60e-02 |
| cellular protein-containing complex assembly | 1.44e-07 | cellular bud neck | 9.04e-03 | mRNA cleavage factor complex | 3.09e-02 | endoplasmic reticulum | 2.00e-02 |
| cytosolic large ribosomal subunit | 2.54e-07 | cytoskeleton organization | 1.27e-02 | peptidyl-lysine modification | 3.53e-02 | plasma membrane | 3.10e-02 |
| mitochondrial translation | 2.87e-06 | protein targeting to mitochondrion | 1.30e-02 | SLIK (SAGA-like) complex | 3.57e-02 |  |  |
| protein binding | 2.29e-05 | transcription initiation from RNA pol II promoter | 1.43e-02 | mating projection tip | 3.95e-02 |  |  |
| mitochondrial large ribosomal subunit | 2.05e-04 | mitochondrial membrane organization | 1.46e-02 | spindle | 3.98e-02 |  |  |
| mitotic sister chromatid segregation | 4.40e-04 | tethering complex | 1.46e-02 | RNA catabolic process | 4.11e-02 |  |  |
| RNA binding | 4.78e-04 | intracellular protein transmembrane transport | 1.49e-02 | spindle pole body | 4.15e-02 |  |  |
| cell division | 5.79e-04 | chromatin binding | 1.55e-02 | DNA-templated transcription, termination | 4.29e-02 |  |  |
| condensed nuclear chromosome kinetochore | 1.01e-03 | intrinsic component of mitochondrial inner membrane | 1.56e-02 | ESCRT III complex | 4.40e-02 |  |  |
| RNA polymerase II, holoenzyme | 1.23e-03 | nuclear migration | 1.90e-02 | dynein complex | 4.58e-02 |  |  |
| cytosolic small ribosomal subunit | 1.31e-03 | RNA pol II transcription factor complex | 2.10e-02 | cortical actin cytoskeleton | 4.66e-02 |  |  |
| SWI/SNF superfamily-type complex | 1.41e-03 | translational initiation | 2.12e-02 |  |  |  |  |
| cytoplasmic microtubule | 2.22e-03 | regulation of gene expression, epigenetic | 2.30e-02 |  |  |  |  |
| protein transmembrane import into intracellular organelle | 2.83e-03 | nucleoside-triphosphatase activity | 2.45e-02 |  |  |  |  |
| transcription elongation from RNA pol II promoter | 3.32e-03 | histone modification | 2.62e-02 |  |  |  |  |
| DNA repair | 6.14e-03 | mRNA metabolic process | 2.62e-02 |  |  |  |  |
| negative regulation of organelle organization | 6.55e-03 | gene silencing | 2.76e-02 |  |  |  |  |
| chromatin remodeling | 7.70e-03 | negative regulation of transcription, DNA-templated | 2.93e-02 |  |  |  |  |

Table S3. Continued.
| Co/S4 |  |  |  | Co/S1 |  | Co/S2 |  |
| --- | --- | --- | --- | --- | --- | --- | --- |
| fungal-type vacuole membrane | 5.90e-10 | aminopeptidase activity | 1.70e-02 | 90S preribosome | 3.00e-11 | cellular bud | 9.60e-03 |
| Golgi membrane | 1.20e-07 | glycogen metabolic process | 1.90e-02 | small-subunit processome | 1.30e-10 | cell periphery | 9.60e-02 |
| vesicle mediated transport | 1.30e-05 | dolichyl-phosphate-mannose-protein mannosyltransferase | 2.20e-02 | endonucleolytic cleavage in 5'-ETS of tricistronic rRNA | 5.00e-06 | Co/S3 |  |
| integral component of plasma membrane | 2.90e-04 | Golgi stack | 2.40e-02 | endonucleolytic cleavage in ITS1 to separate SSU-rRNA | 5.00e-06 | protein refolding | 3.90e-05 |
| protein lipidation | 1.20e-03 | homeostatic process | 2.50e-02 | mRNA binding | 3.20e-05 | antibiotic metabolic process | 3.50e-04 |
| coenzyme binding | 2.70e-03 | porphyrin-containing compound metabolic process | 2.90e-02 | nucleoplasm | 9.70e-05 | cellular response to heat | 8.00e-04 |
| intracellular protein transport | 2.80e-03 | endosome | 3.00e-02 | snoRNA binding | 1.30e-04 | cellular response to chemical stimulus | 8.00e-03 |
| ion binding | 3.00e-03 | glucose binding | 3.30e-02 | endonucleolytic cleavage in ITS1 to separate SSU-rRNA | 3.60e-04 | cytoplasm | 1.30e-02 |
| phospholipid metabolic process | 3.00e-03 | proton-transporting ATPase activity, rotational mechanism | 3.40e-02 | rRNA 2'-O-methylation | 8.10e-03 | cellular catabolic process | 1.60e-02 |
| glycolysis | 4.10e-03 | fatty acid biosynthetic process | 3.40e-02 | box C/D snoRNP complex | 8.90e-03 | intracellular membrane-bounded organelle | 1.80e-02 |
| vacuolar proton-transporting V-type ATPase complex | 4.60e-03 | coenzyme metabolic process | 3.50e-02 | translational termination | 1.10e-02 | chaperone cofactor-dependent protein refolding | 1.90e-02 |
| anion transport | 5.90e-03 | endoplasmic reticulum lumen | 3.70e-02 | cytoplasmic stress granule | 1.30e-02 | organonitrogen compound catabolic process | 2.00e-02 |
| transferring hexosyl group | 6.50e-03 | COPI-coated vesicle | 3.80e-02 | maturation of LSU-rRNA from tricistronic rRNA | 3.70e-02 | unfolded protein binding | 3.90e-02 |
| protein glycolysation | 6.60e-03 | nuclear pore inner ring | 3.80e-02 | preribosome, large subunit precursor | 4.90e-02 | Ras protein signal transduction | 4.60e-02 |
| vacuolar proton-transporting V-type ATPase | 7.60e-03 | vesicle coat | 3.90e-02 |  |  | chaperone binding | 4.60e-02 |
| lipid droplet | 8.20e-03 | trans-Golgi network | 3.90e-02 |  |  |  |  |
| carbohydrate transport | 1.10e-02 | clathrin-coated vesicle | 4.00e-02 |  |  |  |  |
| cellular amino acid biosynthetic process | 1.30e-02 | proteolysis | 4.10e-02 |  |  |  |  |
| ER to Golgi transport vesicle membrane | 1.50e-02 | integral component of endoplasmic reticulum membrane | 8.58e-04 |  |  |  |  |
| secondary active transmembrane activity | 1.60e-02 |  |  |  |  |  |  |

**Table S4.** GO processes for stably expressed genes. Enriched terms were obtained using AMIGO2. FDR score is reported. Biological process terms are indicated in red background, cellular component terms in green and molecular function terms in blue.

| GO-term | FDR |
| --- | --- |
| nucleolus | 6.88e-06 |
| catalytic complex | 2.32e-03 |
| ribosomal large subunit biogenesis | 2.70e-03 |
| preribosome, large subunit precursor | 3.37e-03 |
| cytosolic ribosome | 9.72e-03 |
| ATP binding | 1.34e-02 |
| nucleoplasm | 1.38e-02 |
| phosphate-containing compound metabolic process | 1.87e-02 |
| maturation of 5.8S rRNA | 1.89e-02 |
| cellular protein modification process | 3.42e-02 |
| RNA modification | 3.48e-02 |
| macromolecule methylation | 3.52e-02 |
| ribonucleoprotein complex export from nucleus | 3.83e-02 |
| organonitrogen compound biosynthetic process | 3.96e-02 |
| maturation of SSU-rRNA | 4.07e-02 |
| cleavage involved in rRNA processing | 4.14e-02 |
| ribosome assembly | 4.28e-02 |
| tRNA processing | 4.28e-02 |

**Fig. S1.**
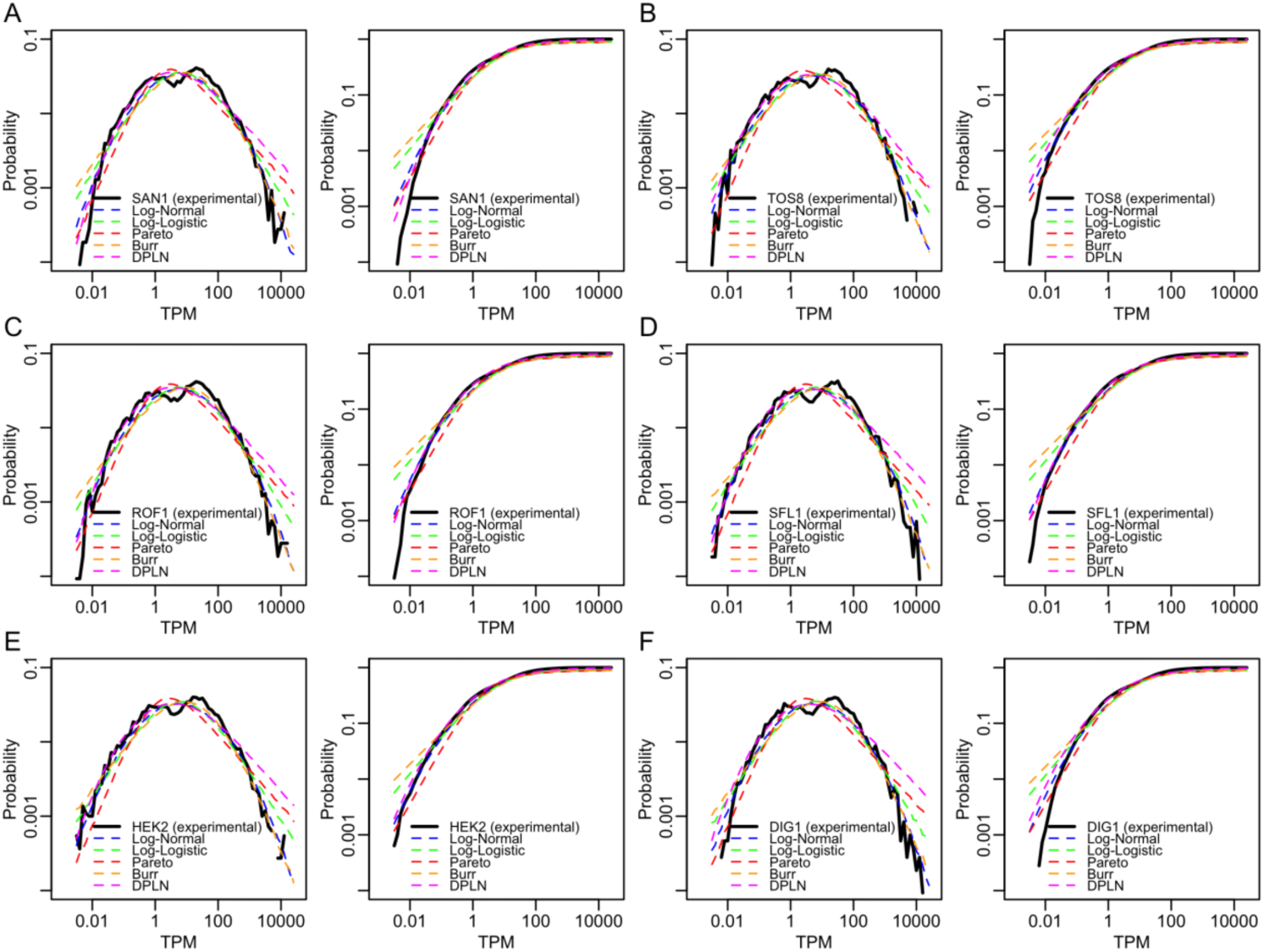
Gene expression distributions (in log scale) against probability density function (PDF, left panels) and cumulative density function (CDF, right panels). Blue, green, red, orange, and magenta curves indicate log-normal, log-logistic, power-law (Pareto), Burr and double Pareto log-normal fitting (DPLN) respectively. The thick black curve corresponds to the experimental distribution in the designated experimental condition (average of 4 replicates): **(A)** SAN1, **(B)** TOS8, **(C)** ROF1, **(D)** SFL1, **(E)** HEK2 and **(F)** DIG1 overexpression. Parameters of the fitted distributions are summarised in Table S1.

**Fig. S2.**
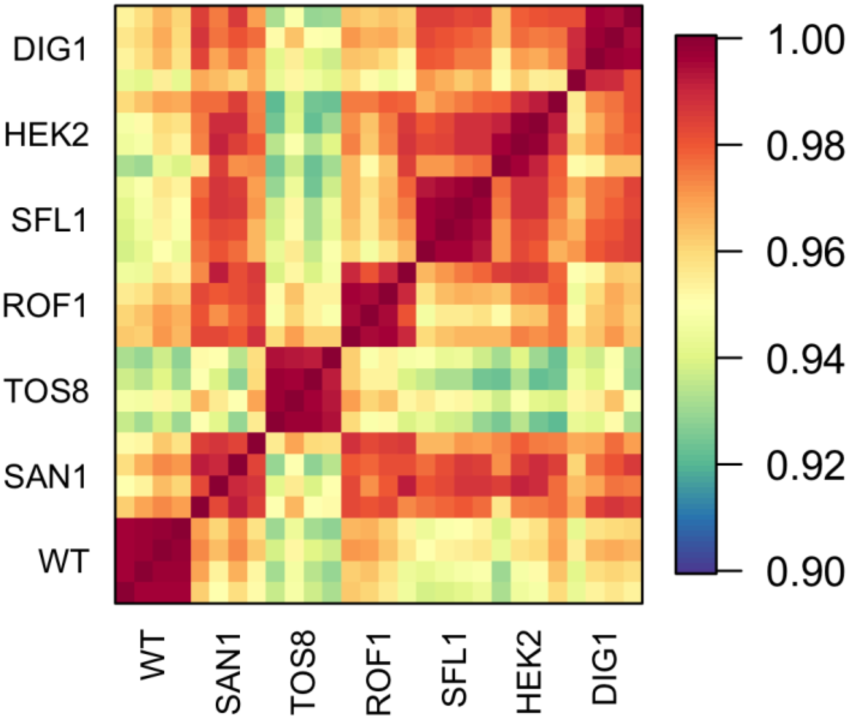
Pearson Correlations between all samples (4 replicates per condition).

**Fig. S3.**
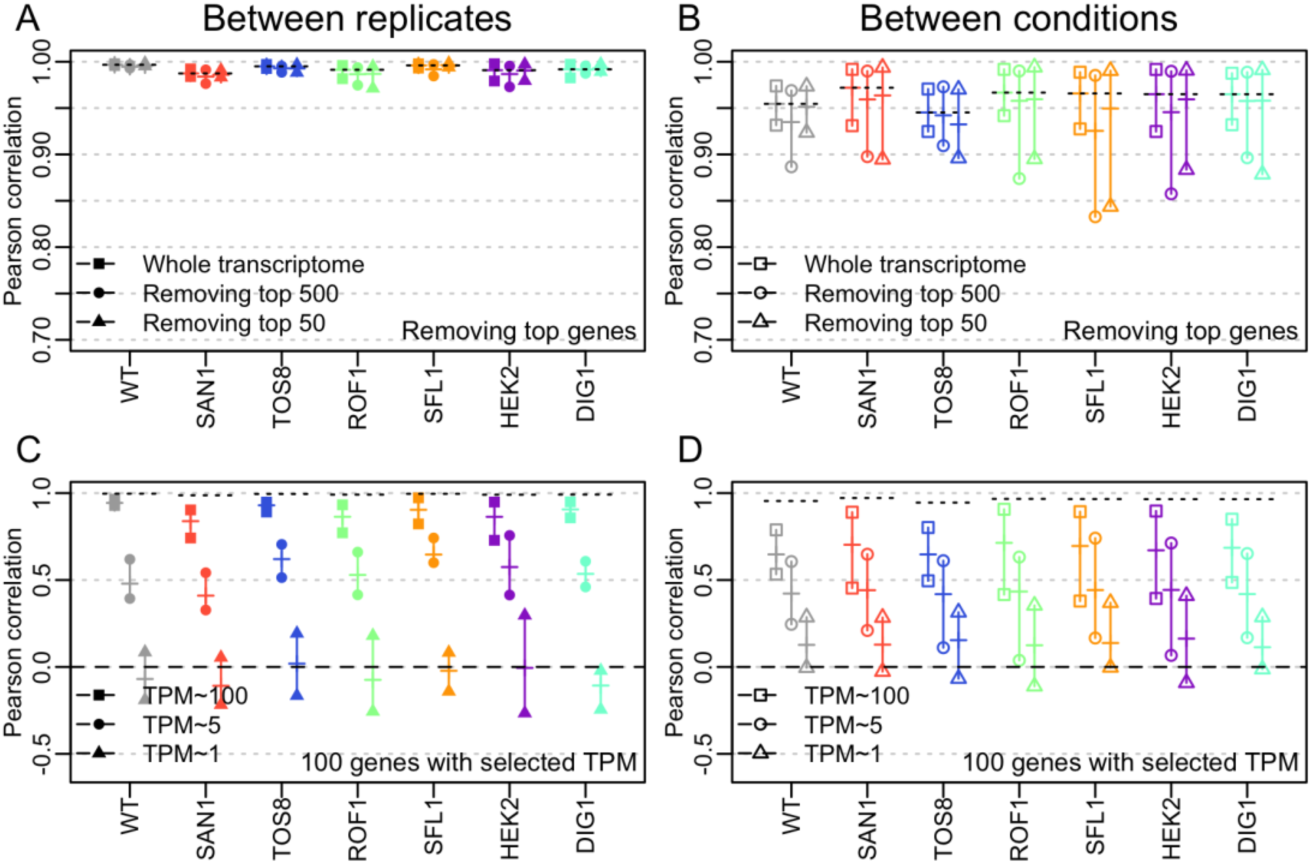
Correlation when **(A,B)** removing top 500 and top 50 most expressed genes in each condition and **(C,D)** selecting 100 genes with TPM ∼ 100, 5, and 1. Filled shapes (on left panels): range (min, max) of values taken by Pearson correlations between replicates (*r*). Empty shapes (on right panels): Pearson correlations against other conditions (*r*_*c*_), for each condition. Dashes: mean correlation values in the range. Dotted lines (repeated in every plot): whole transcriptome average for each condition.

**Fig. S4.**
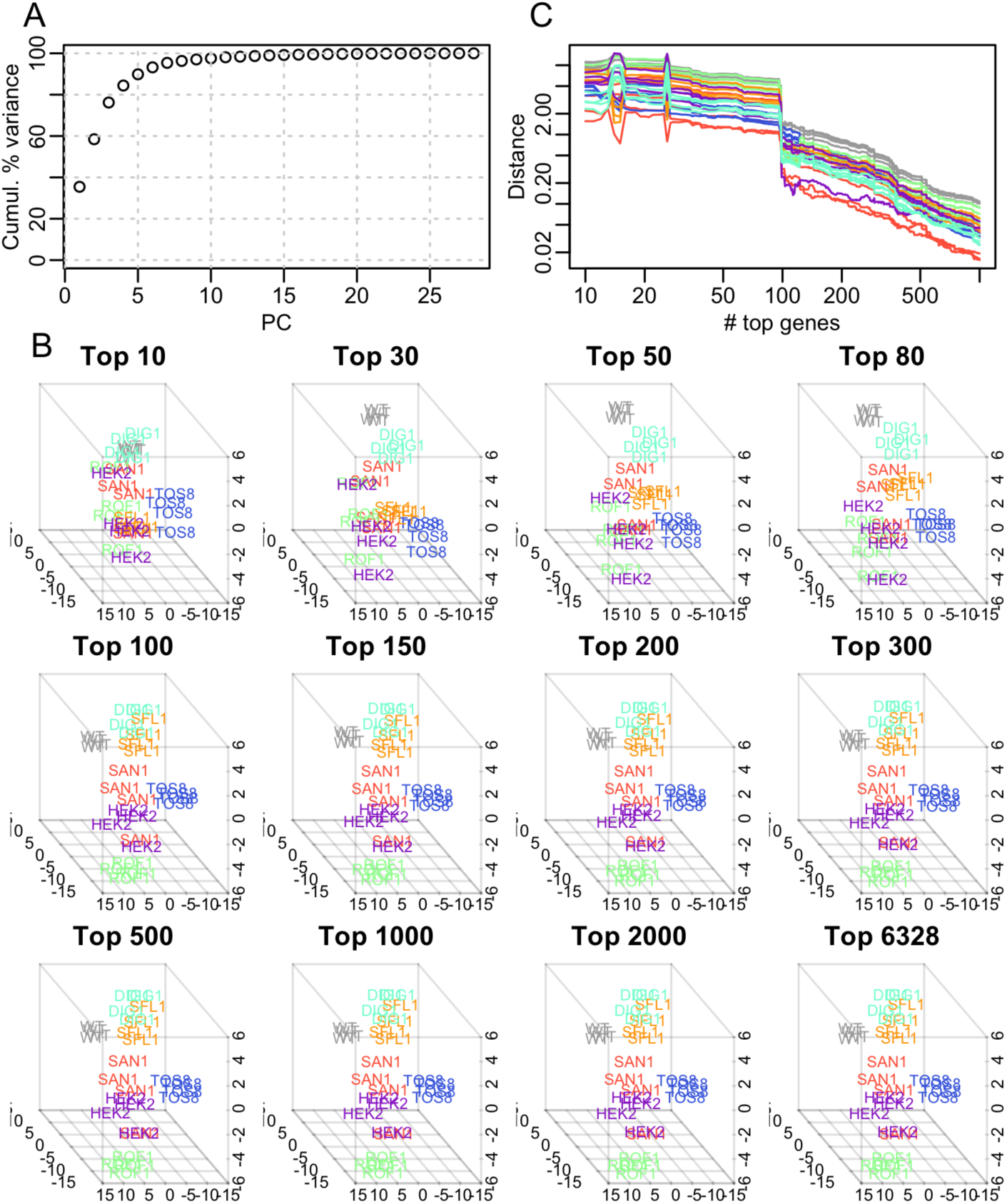
**(A)** Cumulative percentage of variance of the first 28 PCs. **(B)** PCA coordinates (PC1, 2 and 3) of each condition and replicate when selecting top *N* expressed genes with N=10, 30, 50, 80, 100, 150, 200, 300, 500, 1000, 2000 and 6328. **(C)** Euclidean distances between PCA coordinates (PC1, 2 and 3) obtained for *N* = 6,328 and coordinates for *N* = *n* where *n* varies from 1 to 1000. Coordinates stabilise for *N* ≥ 100.

**Fig. S5.**
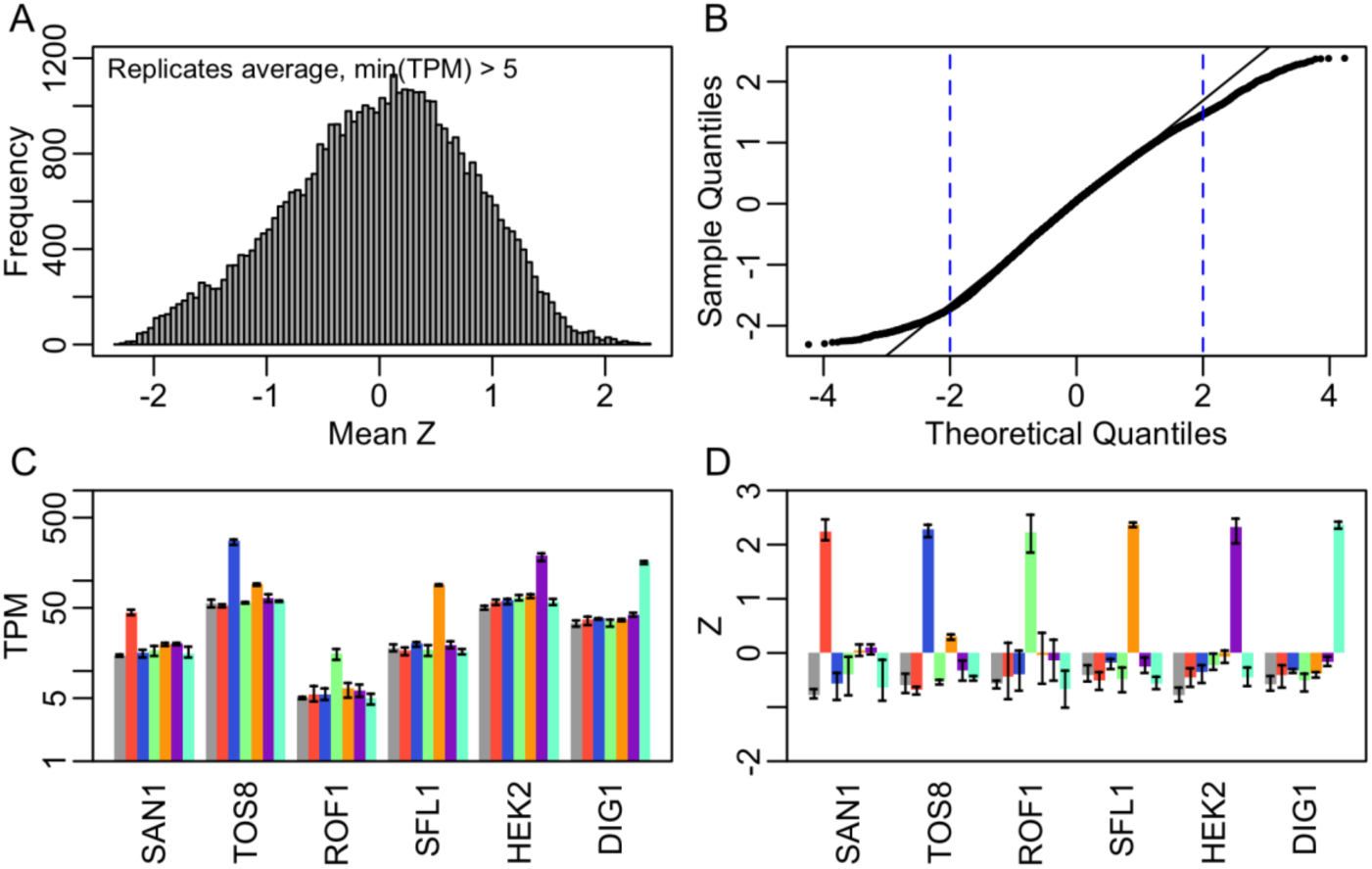
**(A)** Distribution of Z-values averaged per condition, for genes with TPM > 5. **(B)** QQ-plot for testing normality of the distribution of Z-values. Oblique line: normal distribution, thick line: experimental data. Experimental curve follows expected for –2 < *Z* < 2 (dotted blue lines). Mean expression values (TPM) **(C)** and Mean Z-values **(D)** for the 6 overexpressed genes in the 7 experimental conditions. Error bar indicate range of individual replicates.

**Fig. S6.**
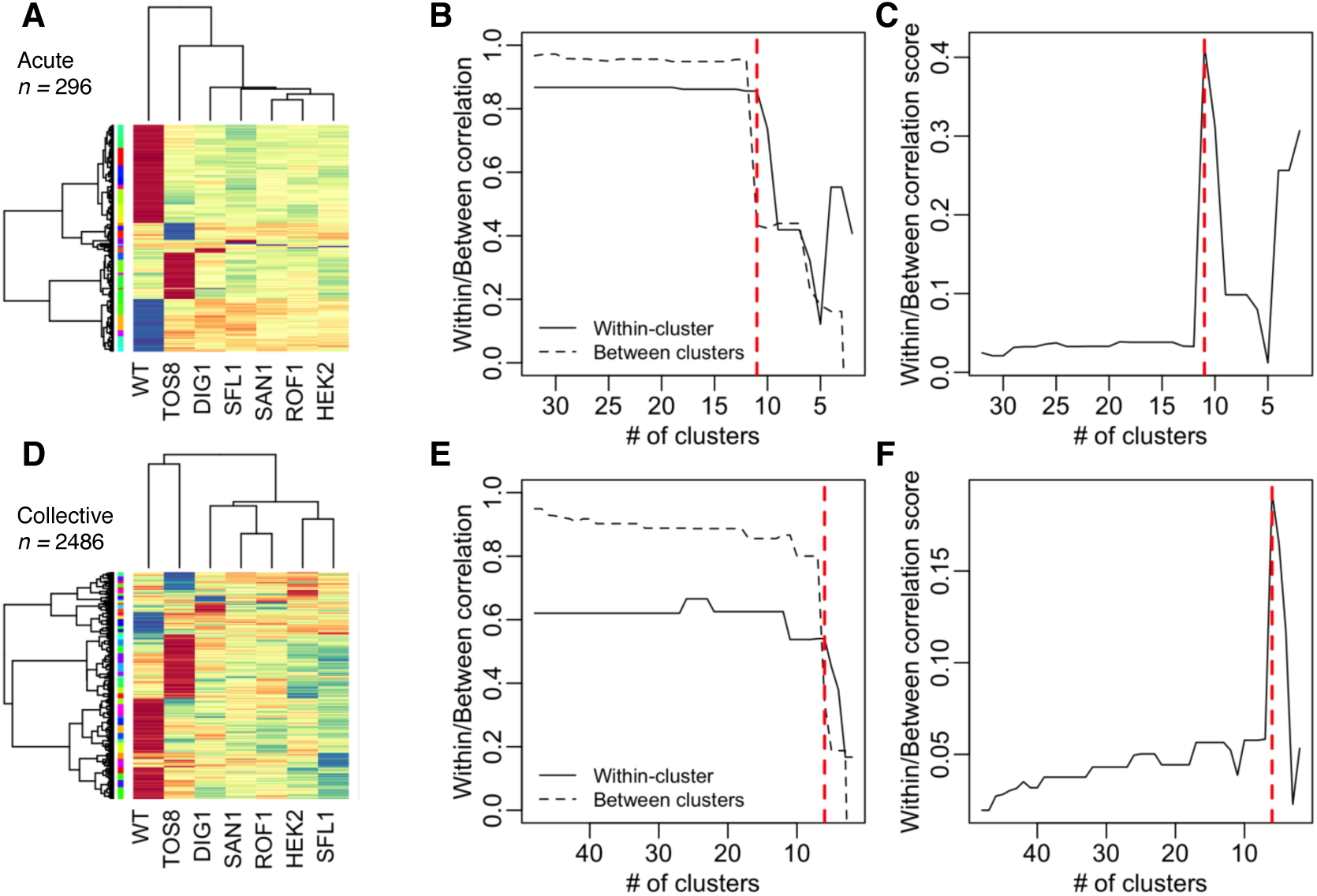
**(A)** The 32 super-clusters obtained by initial hierarchical clustering of the 296 genes forming the acute response. The scale from blue to red indicates up-to down-regulated genes. **(B)** Within-cluster and between clusters correlations when reducing number of clusters. **(C)** Within/Between correlation score when reducing number of clusters. WB-score is obtained such as WB=min(*W*)^2^·(1-max(*B*)), where *W* is the vector of within-clusters correlation, and *B*, the vector of between-clusters. Optimal number of clusters is obtained when WB is maximum. **(D)** Clustering of the 2,486 genes forming the collective response (48 initial super-clusters). **(E)** Within-cluster and between clusters correlations. **(F)** Within/Between correlation score vs. number of clusters.

**Fig. S7.**
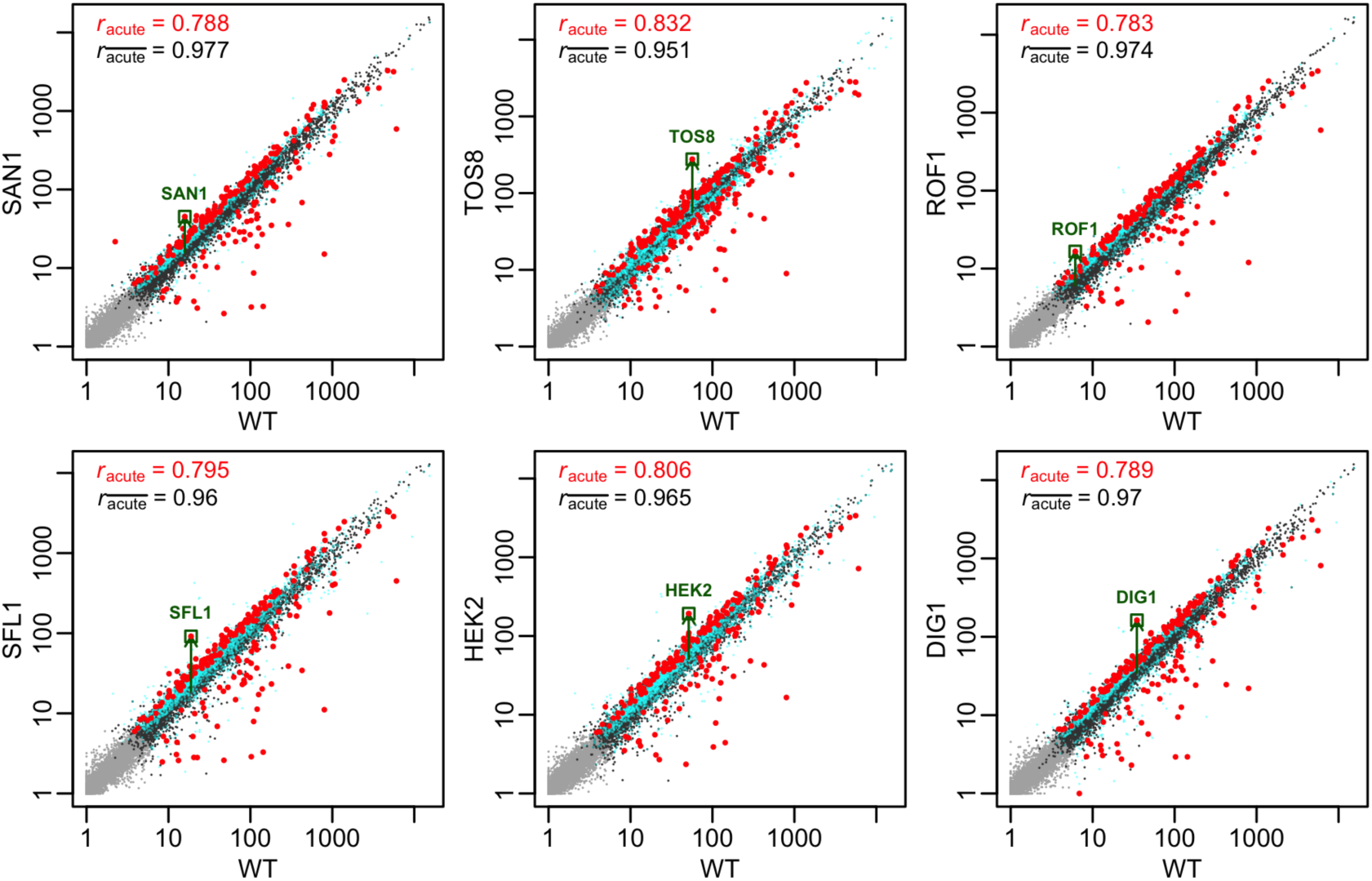
Scatterplot of average expression values (log(TPM+1)) in wildtype vs. other overexpression conditions (average of all replicates per condition). Light grey: non-expressed genes (TPM < 5), dark grey: stably expressed genes (TPM ≥ 5 and |Z| < 1.5), cyan: collective response genes (TPM ≥ 5 and 1.5 ≤ |Z| < 2), red: acute response genes (TPM ≥ 5 and |Z| ≥ 2), dark green square and arrow: overexpressed gene in its respective condition. Correlation for genes from the acute (local) response only *r*_acute_, is displayed in red, and correlation of genes from the transcriptome without acute response, 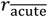, (global structure including collective response genes and stably expressed genes) is displayed in black. Note, since we consider Z-values (normalized expression changes) as a measure for gene response to perturbation, some genes that appear as outlier on the scatterplot (TPM values) may not be considered in the acute response if their expression varies in replicates of the same condition. Also, the overexpressed gene (green square) may not be the most changing gene between wildtype and the respective condition.

**Fig. S8.**
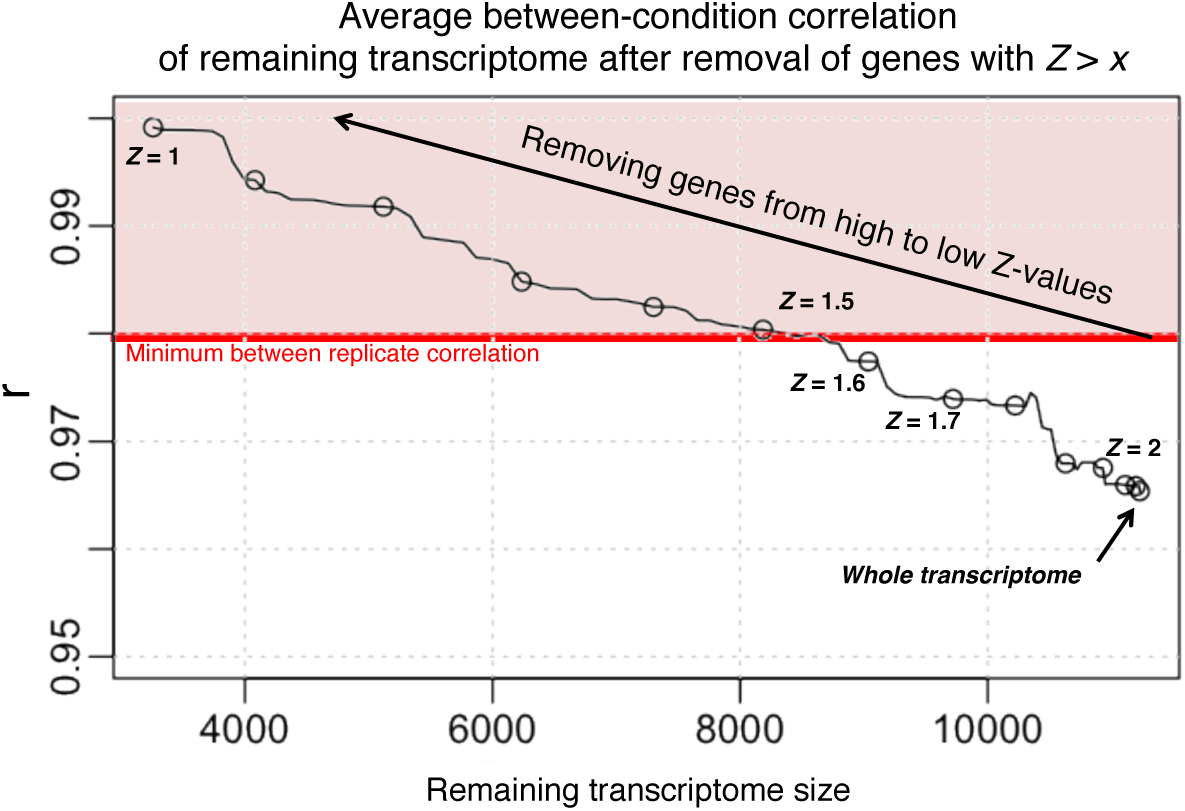
Average cross-correlation values (average of all possible correlations between all replicates of different conditions) when removing genes from high to low absolute Z-values. The whole transcriptome’s (TPM > 0, *n* = 11236) average cross-correlation is located at the utmost right point, indicated by an arrow (*r* = 0.965). Genes from with highest absolute Z-values (max. 2.38) are then removed one by one. The remaining transcriptome size is indicated as *x*-axis. Each dot represents a decrease of 0.1 in the corresponding Z-value. For example, removal of all genes with |*Z*| > 1.7 results in a transcriptome size of *n* = 9719 (x-axis), and cross correlation of *r* ∼ 0.973 (y-axis). The red line indicates the minimum inter-replicate correlation (among all conditions), and is equal to 0.979, which is the maximum shift in correlation value that could be obtained solely from technical noise and biological variability. This threshold value is obtained when removing genes with |*Z*| > 1.5 (approximately, for a transcriptome size, *n* = 8185). Thus, the shift of correlation value from 0.979 (|*Z*| > 1.5) to 0.965 (whole transcriptome) is caused by gene expression changes due to perturbation.

## References

1. Flemming, H. C., Wingender, J., Szewzyk, U., Steinberg, P., Rice, S. A., and Kjelleberg, S.: Biofilms: an emergent form of bacterial life. Nat. Rev. Microbiol. 2016, 14, 563–575.

2. Wang, Z., Gerstein, M., Snyder, M.: “RNA-Seq: a revolutionary tool for transcriptomics”, Nat. Rev. Genet., 2009 Jan, 10, (1), pp. 57–63

3. Albert, R: “Scale-free networks in cell biology”, J. Cell Sci., 2005, 118, pp. 4947–4957.

4. Clauset, A., Shalizi, C. R., Newman, M. E. J.: “Power-Law Distributions in Empirical Data”, SIAM Rev., 51, (4), pp. 661–703

5. Bialek, W: Biophysics: “Searching for Principles” (Princeton University Press, Princeton, 2012)

6. Shannon, C. E.: “A Mathematical Theory of Communication”, Bell System Technical J., 1948, 27, pp. 623–666

7. Rokach, L, Maimon, O.: “Data mining with decision trees: theory and applications” (World Scientific Pub Co Inc., 2008), pp. 81–106

8. Quinlan, J. R.: ‘Induction of Decision Trees. Machine Learning 1’ (Kluwer Academic Publishers, 1986)

9. Soofi, E. S.: “Capturing the intangible concept of information”, J. Am. Stat. Assoc., 1994, 89, pp. 1243–1254

10. Prigogine, I.: “The End of Certainty” (The Free Press, New York, 1997)

11. Kauffman, S.: “At Home in the Universe: The Search for Laws of Self-Organization and Complexity” (Oxford University Press, New York, 1995)

12. Bak, P., Tang, C., Wiesenfeld, K.: “Self-organized criticality: an explanation of 1/f noise”, Phys. Rev. Lett., 1987, 59, (4), pp. 381–384

13. Bak, P., Paczuski, M.: “Complexity, contingency, and criticality”. Proc. Natl. Acad. Sci. U. S. A., 1995, 92, (15), pp. 6689–6696

14. Flemming, H. C., Wingender, J., Szewzyk, U., Steinberg, P., Rice, S. A., Kjelleberg, S.: “Biofilms: an emergent form of bacterial life”, Nat. Rev. Microbiol., 2016 Aug 11, 14, (9), pp. 563–575

15. Koo, H., Allan, R.N., Howlin, R.P.,Stoodley, P.,Hall-Stoodley, L.: “Targeting microbial biofilms: current and prospective therapeutic strategies”, Nat. Rev. Microbiol., 2017 Dec 15 (12), pp. 740–755

16. Selvarajoo, K.: “Order Parameter in Bacterial Biofilm Adaptive Response”, Front. Microbiol., 2018, 9, 1721

17. Tsuchiya, M., Selvarajoo, K., Piras, V., Tomita, M., Giuliani, A.: “Local and global responses in complex gene regulation networks”, Physica A, 2009, 388, pp. 1738–1746

18. Tsuchiya, M., Piras, V., Choi, S., Akira, S., Tomita, M., Giuliani, A., Selvarajoo, K.: “Emergent genome-wide control in wildtype and genetically mutated lipopolysaccarides-stimulated macrophages”, PLoS One, 2009, 4, (3), e4905

19. Piras, V., Tomita, M., Selvarajoo, K.: “Transcriptome-wide variability in single embryonic development cells”, Sci. Rep., 2014, 4, 7137

20. Piras, V., Selvarajoo, K.: “The reduction of gene expression variability from single cells to populations follows simple statistical laws”, Genomics, 2015, 105, pp. 137–144

21. Simeoni, O., Piras, V., Tomita, M., Selvarajoo, K.: “Tracking global gene expression responses in T cell differentiation”, Gene, 2015, 569, pp. 259–266

22. Piras, V., Tomita, M., Selvarajoo, K.: “Is central dogma a global property of cellular information flow?”, Front. Physiol., 2012 Nov 23, 3, 439

23. Felli, N., Cianetti, L., Pelosi, E., Carè, A., Liu, C. G., Calin, G. A., Rossi, S., Peschle, C., Marziali, G., Giuliani, A.: “Hematopoietic differentiation: a coordinated dynamical process towards attractor stable states”, BMC Syst. Biol, 2010 Jun 16, 4, 85

24. Tsuchiya, M., Piras, V., Giuliani, A., Tomita, M., Selvarajoo, K.: “Collective dynamics of specific gene ensembles crucial for neutrophil differentiation: the existence of genome vehicles revealed”, PLoS One, 2010 Aug 11, 5(8), e12116

25. Cromie, G. A., Tan, Z., Hays, M., Sirr, A., Jeffery, E. W., Dudley, A. M.: “Transcriptional Profiling of Biofilm Regulators Identified by an Overexpression Screen in Saccharomyces cerevisiae”, G3 (Bethesda), 2017 Aug 7, 7(8), pp. 2845–2854

26. Cromie, G. A., Tan, Z., Hays, M, Jeffery, E. W., Dudley, M.: “Dissecting Gene Expression Changes Accompanying a Ploidy-Based Phenotypic Switch”, G3 (Bethesda), 2017 Jan 5, 7, (1), pp. 233–246

27. Wagner, G. P., Kin, K., Lynch, V. J.: “Measurement of mRNA abundance using RNA-seq data: RPKM measure is inconsistent among samples”, Theory in biosciences, 2012, 131, (4), pp. 281–285

28. Furusawa, C., Kaneko, K.: “Zipf’s law in gene expression”, Phys. Rev. Lett., 2003 Feb 28, 90, (8), 088102

29. Bengtsson, M., Ståhlberg, A., Rorsman, P.,Kubista, M.: “Gene expression profiling in single cells from the pancreatic islets of Langerhans reveals lognormal distribution of mRNA levels”, Genome Res., 2005 Oct 15, 10), pp. 1388–1392

30. Beal, J.: “Biochemical complexity drives log-normal variation in genetic expression”, IET Engineering Biol., 2017, 1, (1), pp. 55–60

31. Elowitz, M. B., Levine, A. J., Siggia, E. D., Swain, P.S.: “Stochastic gene expression in a single cell”, Science, 2002 Aug 16, 297, (5584), pp. 1183–1186

32. Del Giudice, .M, Bo, S., Grigolon, S., Bosia, C.: “On the role of extrinsic noise in microRNA-mediated bimodal gene expression”, PLoS Comput. Biol., 2018 Apr 17, 14, (4), e1006063

33. Selvarajoo, K.: “Understanding multimodal biological decisions from single cell and population dynamics”, Wiley Interdiscip. Rev. Syst. Biol. Med, 2012 Jul-Aug, 4, (4), pp. 385–399

34. Balsa-Canto, E., Henriques, D., Gábor, A., Banga, JR.: “AMIGO2, a toolbox for dynamic modeling, optimization and control in systems biology”, Bioinformatics, 2016 Nov, 32, (21), pp. 3357–3359

35. Huang, S., Kauffman, S.: “How to escape the cancer attractor: rationale and limitations of multi-target drugs’, Semin. Cancer Biol., 2013 Aug, 23, (4), pp. 270–278

36. Tsuchiya, M., Giuliani, A., Hashimoto, M., Erenpreisa, J., Yoshikawa, K.: “Self-Organizing Global Gene Expression Regulated through Criticality: Mechanism of the Cell-Fate Change”, PLos One, 2016, 11, (12), e0167912

37. Mojtahedi, M., Skupin, A., Zhou, J., Castaño, I. G., Leong-Quong, R. Y., Chang, H., Trachana, K., Giuliani, A., Huang, S.: “Cell Fate Decision as High-Dimensional Critical State Transition”, PLoS Biol., 2016 Dec 27, 14, (2), e2000640

38. Venables, W. N., Ripley, B. D. Modern Applied Statistics with S. (Springer, Fourth edition, 2002)

39. Xiang, Y., Gubian, S., Suomela, B., Hoeng, J.: “Generalized Simulated Annealing for Efficient Global Optimization: the GenSA Package for R”, The R Journal, 2013, 5, (1), pp. 13–28

40. Kleiber, C., Kotz, S.: “Statistical Size Distributions in Economics and Actuarial Sciences” (Wiley, 2003)

41. Reed. W.J., Jorgensen. M.: “The Double Pareto-Lognormal Distribution—A New Parametric Model for Size Distributions”, Commun. Stat. Theory Meth., 2004, 33, (8), pp. 1733–1753

42. Everitt, B. S., Landau, S., Leese, M.: “Cluster Analysis” (Oxford University Press, Inc., New York, 4th Edition, Arnold, London, 2001)

43. Cox, T. F., Cox, M. A.A.: “Multidimensional Scaling” (Chapman and Hall, London, Second edition, 2001)

